# Evaluating Sample Augmentation in Microarray Datasets with Generative Models: A Comparative Pipeline and Insights in Tuberculosis

**DOI:** 10.1101/2021.05.03.442476

**Authors:** Ayushi Gupta, Saad Ahmad, Atharva Sune, Chandan Gupta, Harleen Kaur, Rintu Kutum, Tavpritesh Sethi

## Abstract

High throughput screening technologies have created a fundamental challenge for statistical and machine learning analyses, i.e., the curse of dimensionality. Gene expression data are a quintessential example, high dimensional in variables (Large P) and comparatively much smaller in samples (Small N). However, the large number of variables are not independent. This understanding is reflected in Systems Biology approaches to the transcriptome as a network of coordinated biological functioning or through principal Axes of variation underlying the gene expression. Recent advances in generative deep learning offers a new paradigm to tackle the curse of dimensionality by generating new data from the underlying latent space captured as a deep representation of the observed data. These have led to widespread applications of approaches such as Generative Adversarial Networks (GANs) and Variational Autoencoders (VAEs), especially in domains where millions of data points exist, such as in computer vision and single cell data. Very few studies have focused on generative modeling of bulk transcriptomic data and microarrays, despite being one of the largest types of publicly available biomedical data. Here we review the potential of Generative models in recapitulating and extending biomedical knowledge from microarray data, which may thus limit the potential to yield hundreds of novel biomarkers. Here we review the potential of generative models and conduct a comparative analysis of VAE, GAN and gaussian mixture model (GMM) in a dataset focused on Tuberculosis. We further review whether previously known axes genes can be used as an effective strategy to employ domain knowledge while designing generative models as a means to further reduce biological noise and enhance signals that can be validated by standard enrichment approaches or functional experiments.

## Introduction

Recent advancements in genomics have enabled life scientists to scrutinize the expression of thousands of genes at the same time using next generation sequencing (NGS) [1] and microarray profiling [2]. Unfortunately, the number of samples in such studies becomes a major hurdle, due to a myriad of factors such as small budgets, ethical considerations, or simply because of the low number of subjects available for study [3]. This problem of too few numbers of samples is one of the prominent bottlenecks in biomedical research. A small sample size might not reflect the population well and lead to a decrease in the reproducibility of results [4]. Adding to this, high throughput experiments often accompany significant variations in measured values of transcripts, arising either due to biological variations or experimental noise [5,6]. Therefore, both of these problems have to be addressed while performing any bioinformatics analysis on a genotype or phenotype, for example, identifying candidate gene signatures or the underlying mechanisms for a particular disease.

Many researchers have explored data augmentation techniques for tabular biomedical datasets. Synthetic Minority Oversampling Technique (SMOTE) is a method to overcome imbalanced data problems proposed by Chawla et. Al [7]. The basic concept of SMOTE is to double the number of observations on minor class data to be equivalent to the major class. The way to multiply is by generating synthetic data which is based on the nearest neighbor (k-nearest neighbor) where the nearest neighbor is chosen based on the Euclidean distance between the two data. Even though SMOTE performs well on low-dimensional data it is not effective in the high-dimensional setting, especially in the situations where the signal-to-noise ratio in the data is small.

Machine learning based generative models has been extensively used in a variety of data generating tasks such as in computer vision [8] and natural language processing (NLP) [9]. Generative models have the capability to learn the underlying joint distribution of the data upon training, which can later be used to generate varying numbers of samples based on the learned distribution. Therefore, these are actively used to perform data augmentation during data scarcity to design better classifiers for a classification task.

The most successful of these have been deep generative models such as variational autoencoders (VAEs) and generative adversarial networks (GANs). Apart from dealing with the problem of fewer observations, deep generative models are also able to significantly reduce noise in the datasets thereby improving classification results. Such models have also been used in a variety of problems in biomedical research as well, for example, Beaulieu-Jones BK et al, generated closely resembling biomedical data under privacy restrictions [10]. Jens. et.al. [11] have used GANs for generating binary SNP data from 1000 Genomes Project. Chaudhari et.al. [12] used MG-GAN for improving classification results on small sized cancer datasets. Way et.al. in [13] used variational autoencoders to identify biologically enriched latent space to model cancer transcriptomics. And numerous studies have been performed on in-silico generation of samples for single cell gene expression profiles [12, 14–16].

However, these models have majorly been applied for sequencing based transcriptomic profiles, and not for microarray profiles, which have been an integral part of biomedical research. Experiments based on microarray technology often result in low number of samples, compared to the number of genes available. Since generative models, especially deep generative models require large amounts of data, it’s not clear whether such models can be used for modelling microarray datasets. In addition to this, while high success of these models is evident in the computer vision field, where image datasets also have spatial information embedded, expression data are tabular in nature. Further, GANs may not be able to model the entire data distribution, resulting in mode collapse [17,18] and do not have a robust evaluation method [19,20]. Gaussian Mixture Models (GMMs), one of simplest statistical generative models, do not primarily suffer from mode collapse, and in fact hold the capability to perform at par with GANs with slight modifications [21]. However, only a few studies have used GMMs for modelling biomedical datasets [22,23]. [22] used GMM for identifying gene targets for drug development in mouse and human populations, thus suggesting their application for treatment effects. [23] constructed gene co-expression network from a mixed gene expression datasets for specific tumor subtypes using GMM to deal with extrinsic noise which is not taken into account during correlation studies, which was able to identify novel sub tumor patterns for each tumour. Their applicability for microarray datasets largely remain unexplored.

Hereby, we use tuberculosis microarray dataset, in two settings (Non-Axis and Axis genes [24] settings), for sample augmentation and investigate through statistical and bioinformatics analysis both original as well as synthetic samples generated by various generative models. We use

- Gaussian Mixture Model (GMM) since it models the values from multiple distributions and tends to capture data more efficiently than a single gaussian distribution.
- Variational Autoencoder (VAE), traditionally proposed for input reconstruction in computer vision literature.
- Wasserstein GAN (WGAN) which has been successfully applied to image datasets and is a generalized version of GANs without the general drawbacks of GANs.
- Conditional Tabular GAN (CTGAN) which is a variation of GANs intended to be used for tabular datasets.

To validate the generated gene expression data from each model, we identified differentially expressed genes and associated pathways for both original and synthetic samples and compared them qualitatively. The extra genes and pathways that were found in the synthetic samples and not the original samples were manually validated from existing literature involved in gene expression for TB. This would suggest that the signal-to-noise ratio has been enhanced and consequently new knowledge is discovered. We further perform a thorough comparative analysis across models that have been used in this study.

This paper provides overview of four generative models, namely Gaussian Mixture Model (GMM), and three deep generative models belonging to the class of GANs, Wasserstein GAN (WGAN) and Conditional Tabular GAN (CTGAN), and variational autoencoders (VAEs), followed by evaluation of the datasets in Axis as well as Non-Axis settings.

### Generative Models and Statistical Metrics

#### WGAN

It is a form of generative model that is trained in an adversarial setting of deep neural networks, that is, GAN learns the generative model of a data distribution through adversarial methods. Typically GAN consists of 2 deep neural networks a Generator (G) and a Discriminator (D). The generator takes in an input noise and tries to generate a sample similar to the original dataset, and the discriminator discriminates between the fake and real samples. The two networks are trained simultaneously, as the discriminator improves, the generator learns to generate better and better samples, in the end, a generator that is able to generate samples very similar to the original dataset is obtained.

Let Generator be parameterized by *θ* and the Discriminator by *ϕ*. Let the random input noise (to the generator) be given by z. The sample generated by the generator is given by G_*θ*_(z) and the discriminator output is given by *D_ϕ_*(.). The objective function for the model is given by

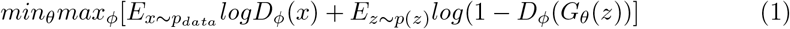

Though GANs traditionally have been proven to perform good at generating samples, they have their own disadvantages. Training of GANs is very unstable and they suffer with other problems like mode collapse, failure to converge, vanishing gradients. To remedy this Wasserstein loss was used. Wasserstein loss is designed to tackle vanishing gradients and mode collapse. To remedy the failure to converge penalizing of discriminator weights is used.

#### CTGAN

Traditionally GANs have been used for synthesizing images, but there have been several extensions that were developed to use GANs on all types of data. Here we explore CTGAN (Conditional GAN for Tabular Data), a GAN based synthesizer developed explicitly for generating tabular data. The problem for tabular generation is formalized as follows

> Let **T** be the table, and it contains **n_c** be the number of continuous variables and **n_d** be the number of discrete variables. Each of the rows represents a vector, say **v**_*i*_. These variables (attributes of the table) have an unknown joint distribution **P**, and each of the row vector **v**_*i*_ is sampled independently from **P**. The aim is to train a model **M**, such that it is able to generate a table **T’**, which follows a similar distribution to **P**.

The preprocessing involves converting discrete variables into one hot encoded variable. For processing continuous variables, a variational Gaussian Mixture Model **(VGM)** is used. The training of the model consists of three parts, **conditional vector, generator loss** and **training-by-sampling**.

The conditional vector is a concatenated vector of all one hot encoded vectors, but with specification of only one category selected (which is to be generated). While training, the models can generate any combination of the one hot vectors (**d**_*i*_), to penalize it for not generating one similar to the input sample (**x_d_*i*_**) is being trained on, by adding cross entropy loss between them. Training by sampling aims to sample efficiently in a way such that all the categories from all discrete variables get sampled evenly during training.

#### VAE

A variational autoencoder generally consists of two networks: a decoder and an encoder. The decoder has a reconstructed output *x*’ for the input *x*, which it aims to be as similar to the input as possible. The encoder-decoder architecture of the model consists of a bottleneck that enforces reduction to the most important features. Using these features the model attempts to approximate the underlying distribution of the input data.

The model contains a decoder function f(·) parametrized by *θ* and an encoder function g(·) parametrized by *ϕ*. The low dimensional latent embedding for an input X is given by *h*=*g_ϕ_(x)* and the reconstructed input *x*’=*f_θ_*(*g_ϕ_(x))*. The aim is to approximate the functions *f* and *g* such that the reconstructed sample *x*’ is similar to the input *x*. To achieve this, a neural network based architecture is used. The decoder function *f* as well as the encoder function *g* are represented by neural networks. The type of layers used in the network depends on the type of data to be generated. The network is then trained to estimate the functions such that the output generated is similar to the input. The loss function for the model is negative log likelihood with a regularizer. The loss function for a sample *x_i_* is given by

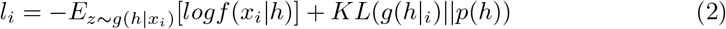

The first term is the reconstruction loss (negative log likelihood). This term encourages the decoder to learn to reconstruct the data. The second term is the regularizer, in this case Kullback-Leibler Divergence. It measures how much information loss occurs when trying to estimate the underlying distribution using the most important features. In VAE *p* is Normal distribution. As the model trains on the input data, it learns to minimize the reconstruction loss, that is the decoder learns to generate as accurate data as possible and minimize the information loss.

#### GMM

A Gaussian Mixture Model (GMM) is a parametric probability density function which is represented as a weighted sum of Gaussian Component distributions. GMMs are commonly used as parametric models to represent continuous measurements. A GMM is given by

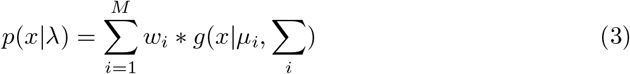

where

x = K-dimensional continuous valued vector
M = GMM component distributions
w_*i*_ = Component Weights
*μ_i_* = mean vector
Σ_*i*_ = covariance vector

Each component distribution (density) *g*(*x*|*μ*,∑) is a K-variate Gaussian distribution of the form

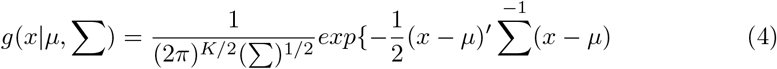

The parameters of the GMM are denoted by λ_*i*_ = {*w_i_, μ_i_*, ∑_*i*_} for i=1….M. These parameters are estimated from training data using the iterative Expectation Maximization method or the Maximum A posteriori estimation from a pre-trained prior model. GMMs have shown excellent performance in clustering tasks, giving better performance than most other clustering algorithm, which can be attributed to the fact that it is an algorithm for density estimation, that is, it is a generative probabilistic model that learns the underlying probability distribution of the data which helps it perform excellently in clustering tasks. In our use case we can use this property of GMM to learn the distribution from training dataset and use it as a generative model to generate synthetic samples.

For our purpose we have used the iterative EM method for estimating the parameters by training the model on available training data.

### Distance metrics to evaluation original vs synthetic data

#### JS Divergence

Jensen-Shannon Divergence or JS Divergence is a method to measure the similarity of two probability distributions. It is based on the Kullback-Leibler Divergence but has some notable and useful differences. The major differences being that it is symmetric and always has a finite value. JS Divergence is a smoothed and symmetrized version of the Kullback-Leibler Divergence Let *P* and *Q* be two distributions 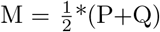, and D(A|||B) represent the Kullback-Leibler divergence between two distributions A and B, then the JS Divergence (JSD) is given by

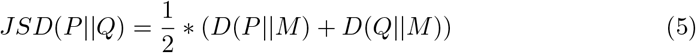

JSD has been applied in bioinformatics, genome comparison, machine learning (to compare generative models).

#### Wasserstein Distance

The Wasserstein Distance or Kantorovich-Rubinstein duality is a distance function which is defined between two probability distributions on a metric space M. It is also called the Earth Mover’s Distance, as informally it can be thought of as the minimum energy cost of transferring a pile of dirt in the shape of one probability distribution to the shape of another distribution. Let P_*r*_ be the real data distribution and P_*g*_ be the generated data distribution the the Wasserstein distance between them is defined as

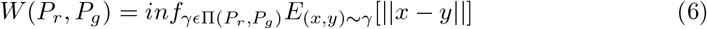

Where Π(P_*r*_, P_*g*_) denotes the set of all joint distributions *γ*(*x,y*) whose marginals are P_*r*_ and P_*g*_ respectively. In intuitive terms γ(*x, y*) denotes the amount of mass being transferred from *x* to *y* in order to transform the distribution P_*r*_ to P_*g*_. The distance then represents the minimum cost to do so.

## Experiments

### Tuberculosis patient data

We select Tuberculosis microarray datasets for sample augmentation since it is a widely studied bacterial infection and numerous studies have been conducted on tuberculosis expression profile [25–28]. For the purpose of selecting datasets, we choose studies with publicly available dataset having both controls and infected samples, more than 100 citations and which were published after 2010. The search queries on pubmed were as follows:

- ((tuberculosis) AND (expression) AND ((case) OR (study)))
- ((tuberculosis) AND (expression) AND (dataset))
- (tuberculosis) AND (microarray)
- ((tuberculosis) AND (gene) AND (signature))

GSE37250 was chosen to be the dataset of choice because of relatively large sample size and having clear distinction between samples having latent TB infection and active TB infection.

The complete normalized dataset has 32 distinct chromosomes, and a total of 47323 genes and 537 adult patients (Cape Town, South Africa (n=300) and Karonga, Malawi (n=237) who were either HIV+ or HIV - with either active TB, LTBI or OD).

There are 6 distinct classes amongst which we use 2 classes as follows:

- Control Group [83 patients] → Group 1
- Active TB, No HIV [97 patients] → Group 2

After collection of Data, preprocessing was done and the steps followed for the preprocessing are as follows:

- 47323 ILLUMINA IDs were converted to 47323 gene names and then the duplicate genes were removed by taking the mean of the gene expression data. Final number of unique genes is 34602.
- Adding 1 and the minimum value of the dataset to all other values.
- Log2 transformation, if D represents the original dataset then D=log2(D+min(D)+1).

The Log2 transformed data was passed onto these models, where feature selection using PCA was done to:

- Shorten the feature space from 34602 genes to a computationally economical space consisting of around 100 features.
- Preserve 99% of the variance in the dataset.

However, PCA was not performed for the data consisting of Axes genes since the feature space was small.

### Whole gene set and axes genes

In order to check the capacity of generative models to reduce noise while retaining as many biologically significant characteristics as possible, we train and evaluate our models under two different settings - whole gene set (Non Axes setting) and axes genes set (Axes setting). Axes genes [24] are a set of axes with highly conserved variation. For our studies, we retain the axis 7 genes among the nine axes (167 in number), which is specified for viral infection.

Under Non Axes settings, since the number of genes is extremely high which may result in over-fitting, principal component analysis (PCA) is performed to induce dimension reduction, and principal components responsible for 99% variation are retained for further analysis.

### Training of Generative Models

For our purpose the data is first split into 2 different sets (for 2 different groups active TB and healthy) and then each set is passed on to the model (thus 2 different models) as input. Hyper-parameter tuning is performed to select the best parameters for each generative model under both the settings. For WGAN it was observed that shallow neural networks showed better performance overall than deeper networks based on our evaluation criteria, hence each encoder (generator) and decoder (discriminator/critic) sub-network was designed with only two to three latent layers. The same number of hidden layers for encoder and critic are used, with all the layers having leaky relu activations except the final critic layer, which has no activation. We also used a gradient penalty which has shown to be a better variant of WGAN. The input to the model is a vector of gene expression values for all the genes for a given patient. For both VAE and WGAN, the optimizing function used is Adam Optimizer and the learning rate is 5e-4. For CTGAN, the pip package provides the tuning of a few hyper-parameters passed as arguments. Similarly for GMM, the scikit-learn package was used where AIC and BIC for the fitted models was plotted (Fig. 1) in axes and non axes genes settings respectively. Optimal number of components (5 in each case) were selected from the resulting curve where a dip was reported.

**Figure 1.**
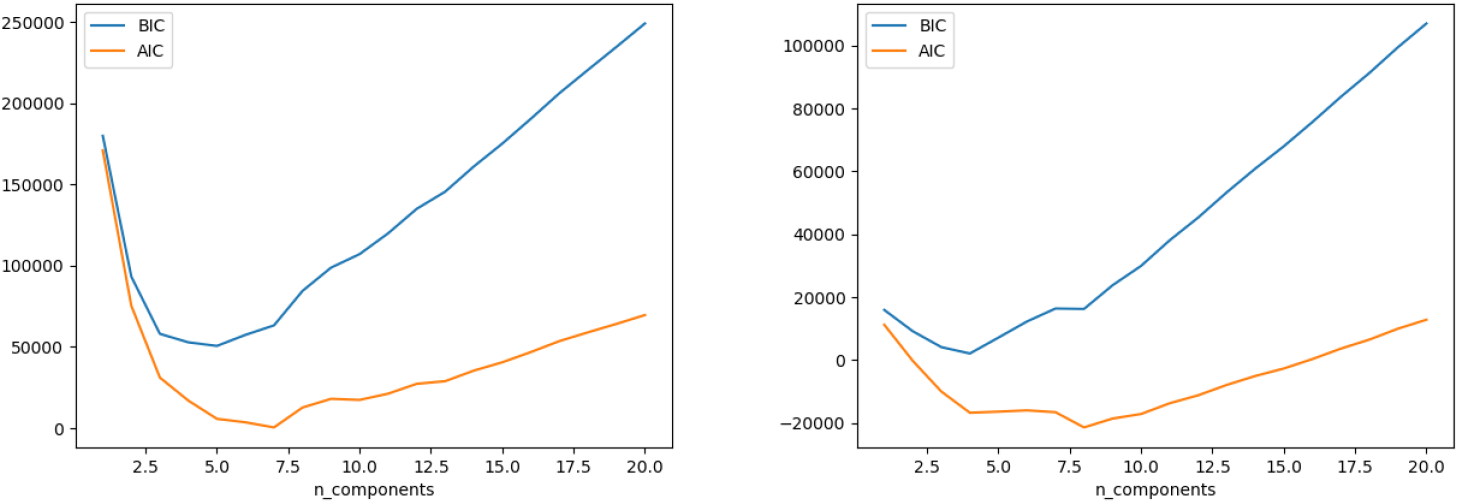
AIC-BIC curve for TB class. Non Axes (Left) and Axes (Right).

For the deep generative models, PyTorch framework, a popular deep learning library in python, was used to train and generate results.

Under both Axis and Non-Axis settings, two different generative models were trained for tuberculosis and control groups.

### Evaluation of generative models

To assess the generative power of each model, a comparative statistical as well as bioinformatics analysis was done on both original as well as synthetically generated samples from each model, for both the settings. t-SNE plots between generated and original data in both settings have been plotted, JS Divergence and Wasserstein Distance plotted against increasing Epochs (in case of DL models), increasing Number of mixture components (in case of GMM) and increasing number of training samples have also been reported in statistical analysis.

Differentially expressed (DE) genes were calculated for each of the datasets by using t-tests between the case and the control. Log2(Fold Change) cutoff was set to 0.5 and adjusted p-value (holm correction) cutoff was set to 0.05. Overlap of DE genes was calculated between generated data and original data. Analysis of genes overlap was also performed with KEGG disease pathway (Pathway: hsa05152) genes for Tuberculosis.

DE genes were identified and consequently pathway analysis was performed on a larger number of generated samples than present in the original data (550 samples). The number of overlapping and extra DE genes in synthetic DE genes set as well as pathways were compared with the original DE genes set in each case. Additionally, to evaluate the extra information obtained from synthetic data, the DE genes set and the associated ontologies/pathways were compared with known literature around Tuberculosis. Majorly, two references [25,26] describing the expression profile of Tuberculosis were used. While it is expected that the generative models should be able to capture the underlying biological characteristics of the two genotypes with reduced signal to noise ratio, we further hypothesized that the synthetic DE genes set may increase the enrichment in each pathway comparatively due to extra genes.

## Evaluation

### Analysis of the generated data using t-SNE plots

Comparison of high dimensional data can be performed using t-SNE which maintains the high dimensional relationships and can plot the data in lower dimensional space. Our qualitative assessment using these plots (Fig. 2, 3, 4, 5) reveal the patterns captured by these models. Data generated using GMM shows a very close relationship with the original data in all the scenarios, thus showing potential for use in scenarios where data needs to be precisely replicated. WGAN and CTGAN show deviations from the original data which indicates that these models may be learning signals not present in the original data. This behaviour leads to extra biological information which is explored in the subsequent sections.

**Figure 2.**
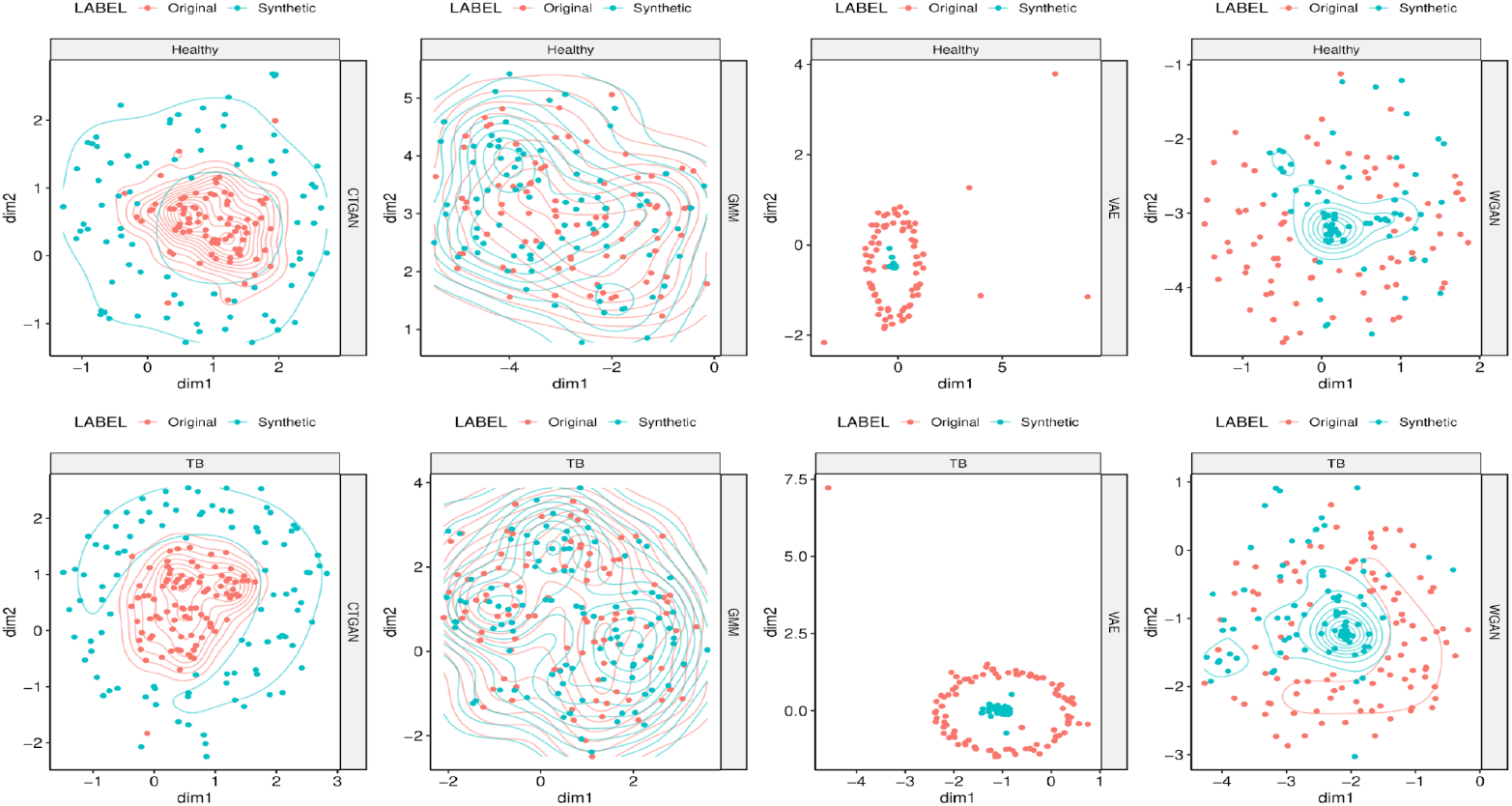
t-SNE plots for generative models comparing generated data to original data for Healthy and TB samples.

**Figure 3.**
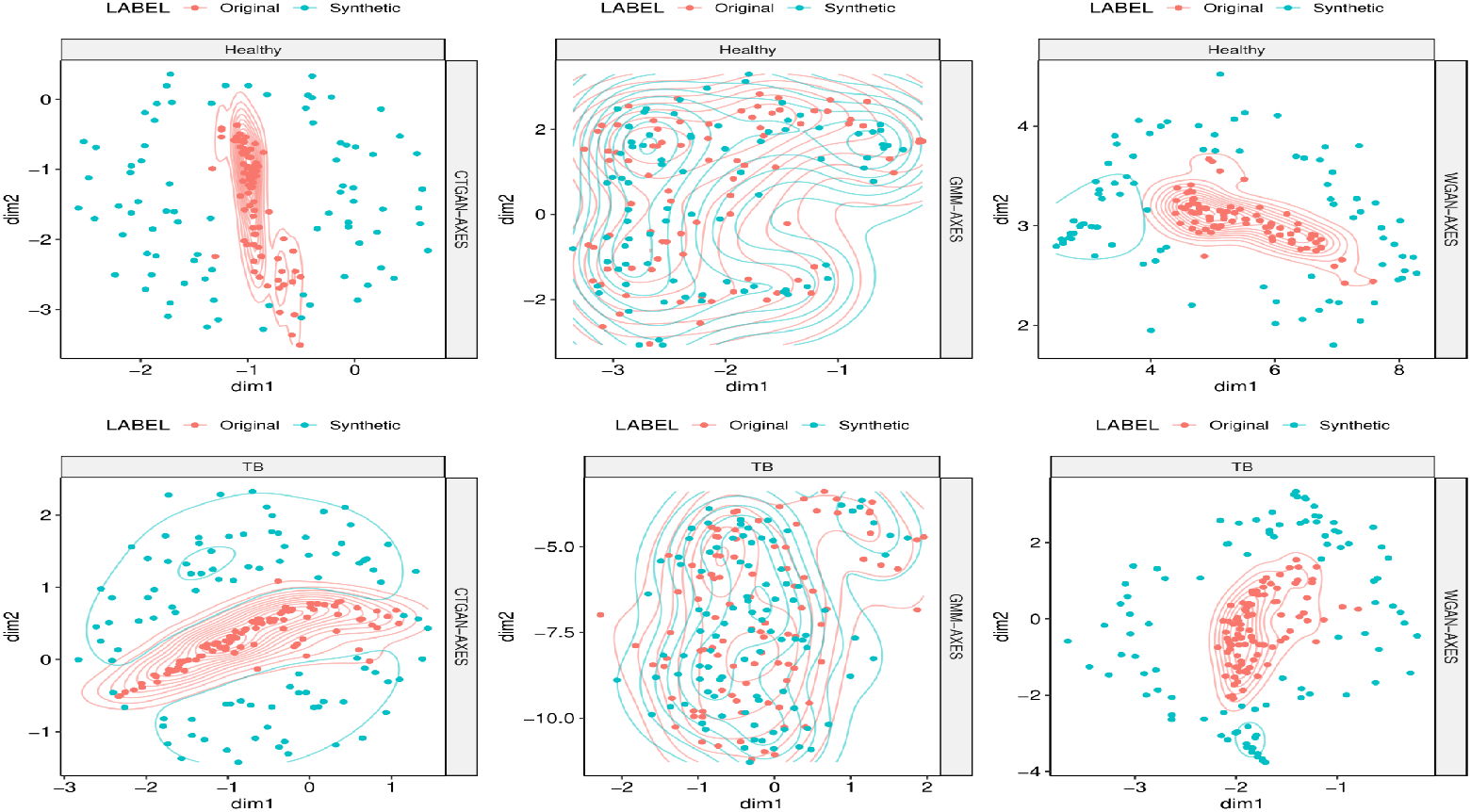
t-SNE plots for generative models comparing generated data to original data for Healthy and TB samples under Axes Genes setting.

**Figure 4.**
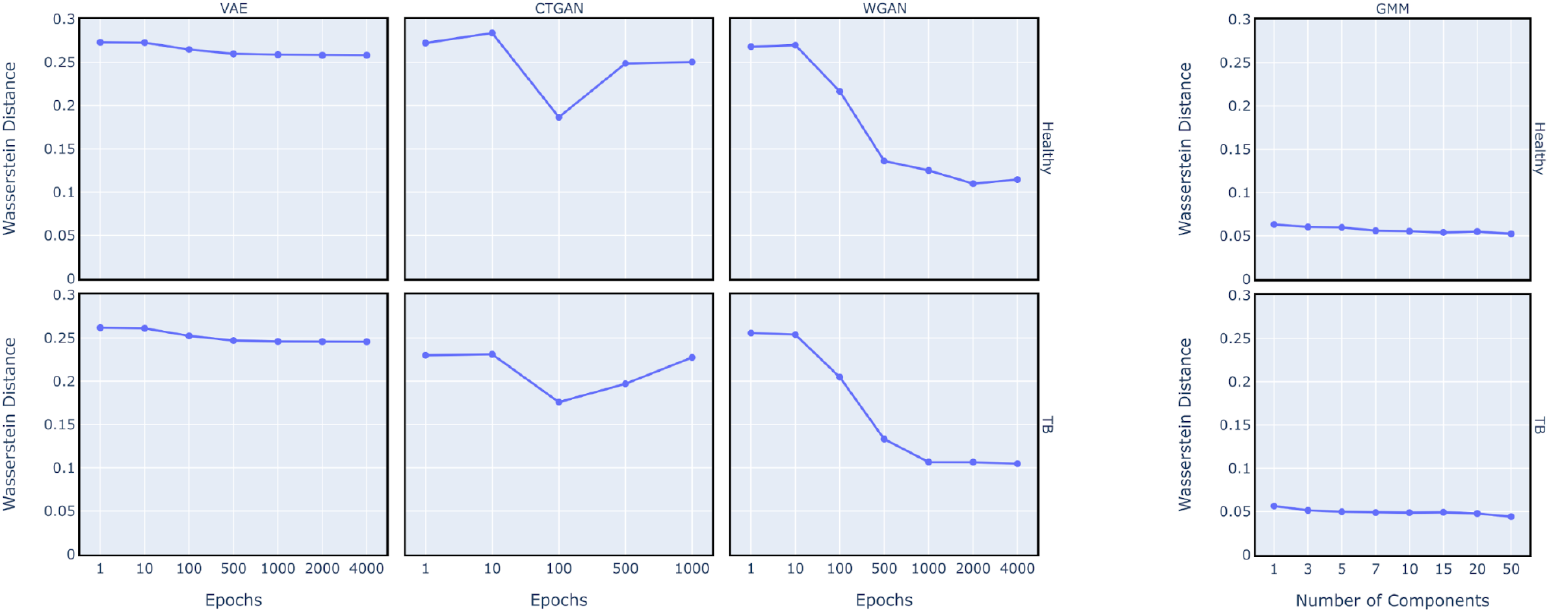
Epoch-wise Wasserstein distance for the three DL models and Number of components wise Wasserstein distance for GMM.

**Figure 5.**
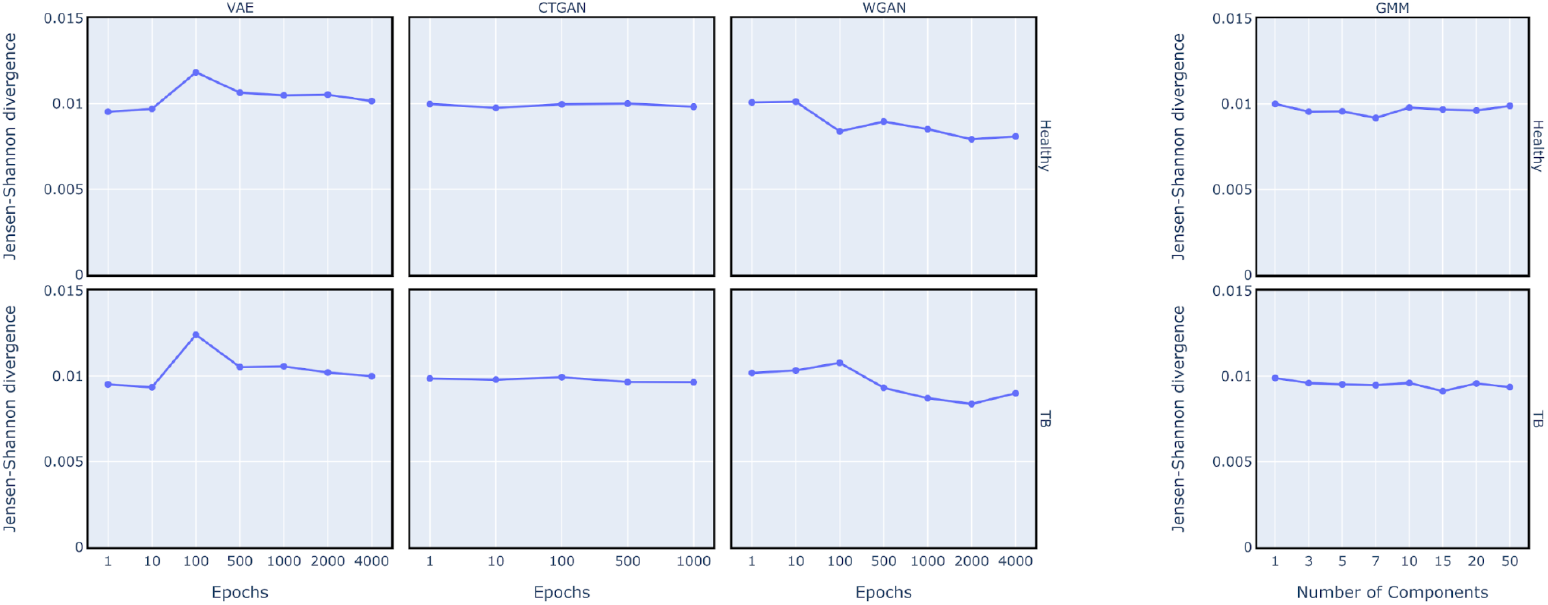
Epoch-wise JS divergence for the three DL models and Number of components wise JS divergence for GMM.

### Comparison of models using Wasserstein distance and JS Divergence

To quantitatively test the learning of models and similarity of generated data, epochwise Wasserstein distance and Jensen-Shannon divergence between generated and original distribution was plotted for WGAN, CTGAN and VAE. Similarly for GMM, both of these metrics were calculated for a set of number of mixture components. It was observed that for WGAN both the metrics showed regular decrement in the values barring a few increments with increasing number of epochs, regular decrement in Wasserstein distance with epochs is expected since WGAN uses this distance as its loss function. CTGAN shows no particular trend in either of the metrics. We also observe similar or lower values for GMM on both metrics indicating that the generated data closely resembles the original data. A slight tilt is observed when increasing the number of mixture components for GMM, which is also expected since larger number of mixture components tend to overfit the training data.

### Comparison of models on number of training samples using the two metrics

It has been hypothesized that generative models generally give better results when trained on larger training sample size. To test the same, we randomly sampled n patients from the data consisting of 83 and 97 samples in healthy and TB classes respectively (where n takes discrete values between 25 and 80 (Fig. 6)). Using Wasserstein distance as the metric we see a trend of decreasing values on increasing training sample size in all the models except CTGAN where an upward trend is observed. However, when observing JS divergence, a clear trend is not observed in CTGAN, WGAN and GMM. VAE shows a general decreasing trend albeit with a few irregularities. These two plots indicate that CTGAN might not be learning the distribution of the dataset as well as the other models.

**Figure 6.**
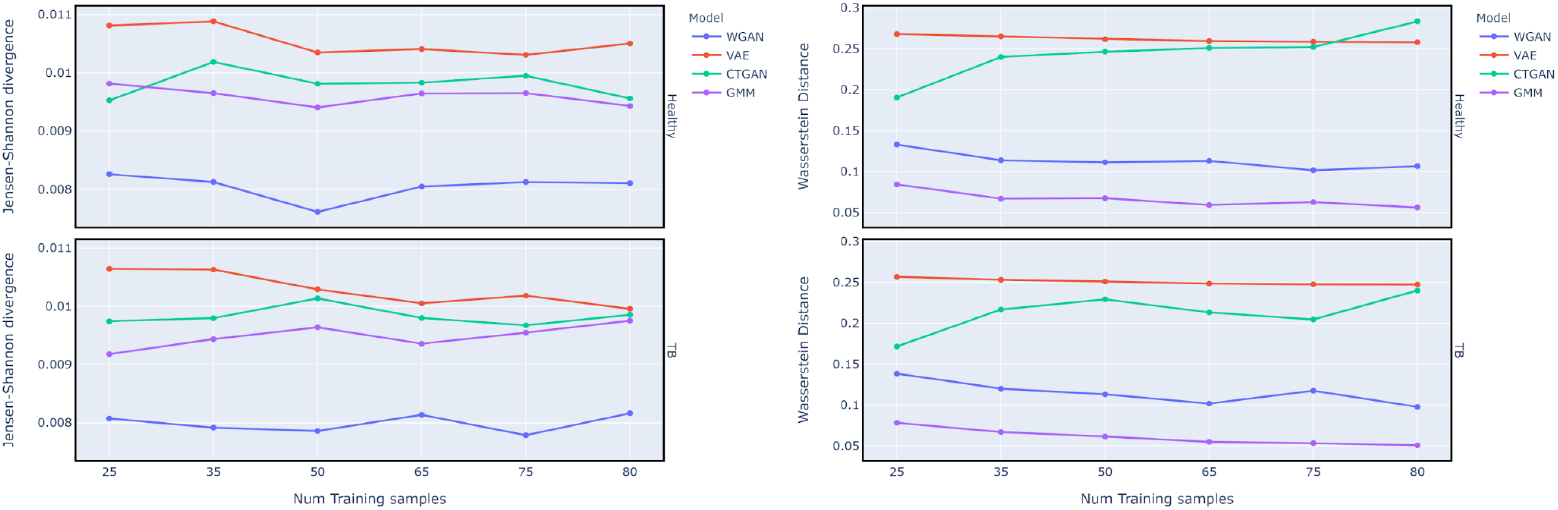
Variation of Wasserstein distance (right) and JS Divergence (left) between generated and training samples with increasing training sample size.

### Analysis of differentially expressed genes from the generated models

In this section, we analyse the differentially expressed (DE) genes sets for generated datasets and compare the overlap with the original and KEGG genes sets. The KEGG gene set for tuberculosis provides 179 genes which have been found to be biologically significant in the infection. The original dataset (Non-Axes setting) gives a list of 121 DE Genes out of which only 4 match with the KEGG gene set.

Table 2 shows that with increasing number of training samples, the overlap of DE genes with original data improves but the same cannot be conclusively said for overlap with KEGG (hsa05152) genes. However, a larger overlap of genes with KEGG genes was indeed observed in the generated datasets compared to the original data. Large number of extra DE Genes and a generally low overlap of DE genes with the original dataset indicates that CTGAN is not learning from the data as well as the other models.

**Table 1.** The datasets which matched our inclusion criteria were:

| GEO Accession ID | Year | Number of Samples |
| --- | --- | --- |
| GSE56153 | 2014 | 36 |
| GSE37250 | 2013 | 180 |

**Table 2.** Overlap of DE Genes from models with original data and KEGG genes observed with increasing training sample size.

| Model | Training sample size | Overlap with KEGG genes | Number of DE Genes | Overlap with DE Genes from original data |
| --- | --- | --- | --- | --- |
| CTGAN | 25 | 3 | 398 | 23 |
| CTGAN | 35 | 7 | 791 | 75 |
| CTGAN | 50 | 14 | 1436 | 72 |
| CTGAN | 65 | 1 | 180 | 29 |
| CTGAN | 75 | 8 | 211 | 12 |
| CTGAN | 80 | 1 | 411 | 26 |
| GMM | 25 | 4 | 226 | 68 |
| GMM | 35 | 2 | 138 | 68 |
| GMM | 50 | 5 | 180 | 94 |
| GMM | 65 | 4 | 204 | 104 |
| GMM | 75 | 4 | 164 | 96 |
| GMM | 80 | 6 | 172 | 111 |
| VAE | 25 | 8 | 325 | 96 |
| VAE | 35 | 8 | 227 | 100 |
| VAE | 50 | 8 | 222 | 110 |
| VAE | 65 | 7 | 272 | 116 |
| VAE | 75 | 5 | 243 | 109 |
| VAE | 80 | 6 | 244 | 115 |
| WGAN | 25 | 7 | 420 | 97 |
| WGAN | 35 | 11 | 911 | 93 |
| WGAN | 50 | 8 | 295 | 84 |
| WGAN | 65 | 12 | 957 | 87 |
| WGAN | 75 | 6 | 454 | 102 |
| WGAN | 80 | 11 | 517 | 121 |

Table 3 compares the overlap of DE genes with the original and original axes data in Non-Axes and Axes settings respectively with an increasing number of generated samples.

**Table 3.** Overlap of DE Genes with Original in Axes and Non-Axes settings.

| Original Data (Non-Axes setting),<br>Original Data (Axes setting) | 121 DE Genes, 34 Axes DE Genes |
| --- | --- |
| GMM | 430 DEG with an overlap of 118 |
| WGAN | 862 DEG with an overlap of 120 |
| CTGAN | 520 DEG with an overlap of 15 |
| GMM (250 Samples) | 261 DEG with an overlap of 120 |
| WGAN (250 Samples) | 863 DEG with an overlap of 120 |
| CTGAN (250 Samples) | 1191 DEG with an overlap of 54 |
| GMM (550 Samples) | 183 DEG with an overlap of 109 |
| WGAN (550 Samples) | 684 DEG with an overlap of 120 |
| CTGAN (550 Samples) | 1691 DEG with an overlap of 80 |
| GMM (Axes setting) | 18 Axes DEGs with an overlap of 18 |
| WGAN (Axes setting) | 18 Axes DEGs with an overlap of 10 |
| CTGAN (Axes setting) | 32 Axes DEGs with an overlap of 15 |
| GMM (250 Samples in Axes setting) | 28 Axes DEGs with an overlap of 26 |
| WGAN (250 Samples in Axes setting) | 74 Axes DEGs with an overlap of 31 |
| CTGAN (250 Samples in Axes setting) | 52 Axes DEGs with an overlap of 24 |
| GMM (550 Samples in Axes setting) | 39 Axes DEGs with an overlap of 34 |
| WGAN (550 Samples in Axes setting) | 107 Axes DEGs with and overlap of 32 |
| CTGAN (550 Samples in Axes setting) | 50 Axes DEGs with an overlap of 25 |

With an increase in sample size, the DE Genes overlap was found to be increasing in case of CTGAN. A lot of extra DE Genes were reported with large sample sizes. Similar to the Non-Axes settings, a clear increase in both, the overlap and the number of DEGs is to be seen under Axes setting. The extra DE genes obtained in this section are explored in the subsequent section.

### Information discovery by comparing the Molecular signals associated with Tuberculosis

It was observed that the models generate extra differentially expressed genes when compared to original data. To explore the significance of these genes we compare our model’s r1esults with published literature about Tuberculosis expression [25, 26].

The authors in [26] have curated the list of differentially expressed genes from 16 different Tuberculosis datasets and composed the genes in components which are presented as a cartoon of the 380-gene-meta-signature in figure 1.e of the paper [26].

We assess if a generative model manages to capture DE Genes which are part of biological components present in figure 1.e of the paper [26]. but are not differentially expressed in the original dataset.

#### Non Axes setting

All the DE Genes present in the original dataset were also found in the list of DE Genes from all the 3 generative models. WGAN, GMM and CTGAN were observed to differentially express 11, 8 and 15 extra genes respectively which were not present in the original dataset (Table 4). Of these extra genes, a majority were associated with the interferon inducible category. These genes are simulated by interferons and are also known as ISGs (interferon stimulated genes) that take part in activating host’s antiviral immunity [29]. It has been observed that the ISGs (interferon stimulated genes) from IFN type 1 pathway antagonise the signalling downstream of IFN-γ [30]. The upregulation of these ISGs is of relevance in Tuberculosis exacerbation. The generative models seem to be able to capture this relationship as many of the extra differentially expressed genes belong to these upregulated ISGs. With CTGAN, we observe most number of extra genes, of these 2 (STAT1, JAK1) are part of the IFN-γ signalling pathway which has been described to be of paramount importance in Tuberculosis [26]. The presence of STAT1, JAK1 and CAMP in CTGAN is interesting due to the fact that none of the genes from this category were present in either original data or either of WGAN or GMM, this is in contrast to WGAN and GMM where all the categories associated with extra genes had at least one gene present in the list of DE Genes from Original data. We also observe the occurrence of 3 moderately expressed genes (JUN, TLR2, JAK1) in CTGAN, this occurrence could be due to the fact that CTGAN gives the most number of differentially expressed genes on the same cutoff as different models and has a significantly different t-SNE plot compared to the original data. WGAN and CTGAN give 2 and 3 extra DE Genes respectively from the Complement and granules category. This category has been shown to be of importance to *myobacterium tuberculosis* [31–33].

**Table 4.**
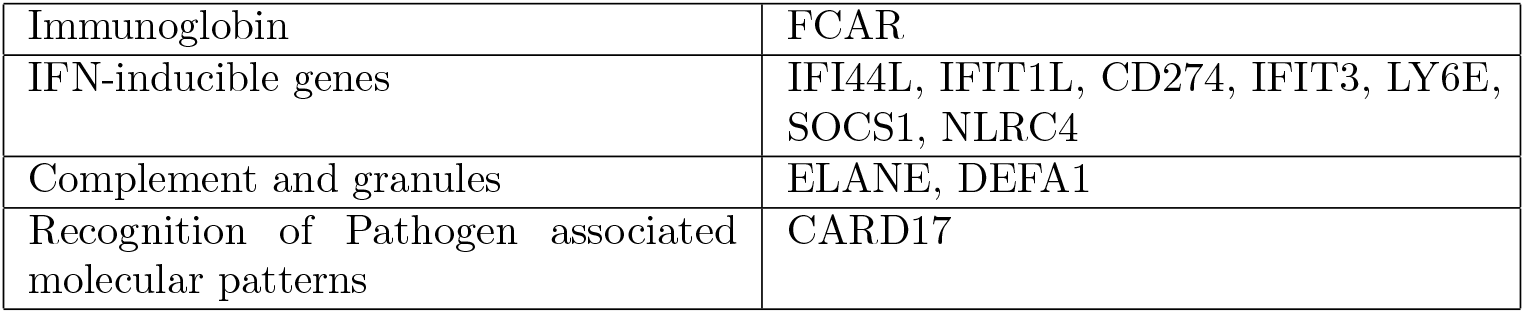
Components associated with extra DE Genes by WGAN with 550 sample.

**Table 5.** Components associated with extra DE Genes by GMM with 550 sample.

|  |  |
| --- | --- |
| IFN-inducible genes | RSAD2, SOCS1, LY6E, IFITM3, IFIT3, IFI44 |
| Complement and granules | DEFA1 |
| Recognition of Pathogen associated molecular patterns | CARD17 |

**Table 6.** Components associated with extra DE Genes by CTGAN with 550 sample.

|  |  |
| --- | --- |
| IFN signalling | STAT1, JAK1 |
| IFN-inducible genes | IFI44, IFIT1, LY6E, RSAD2, OAS1, NLRC4 |
| Pro-inflammatory | CAMP |
| Complement and granules | ELANE, CR1, DEFA1 |
| Recognition of Pathogen associated molecular patterns | TLR4, TLR2, JUN |

**Table 7.** Components associated with extra DE Genes by WGAN Axes setting with 550 samples.

|  |  |
| --- | --- |
| Recognition of Pathogen associated molecular patterns | CASP5 |
| IFN-inducible genes | RSAD2, IFIT1, IFI35, CMPK2 |
| IFN-signalling | STAT1 |
| Genes present in Original but not DE Genes from WGAN | None |

**Table 8.** Components associated with extra DE Genes by GMM Axes setting with 550 samples.

|  |  |
| --- | --- |
| IFN-inducible genes | RSAD2 |
| IFN-signalling | STAT1 |
| Genes present in Original but not in DE genes from GMM | None |

**Table 9.**
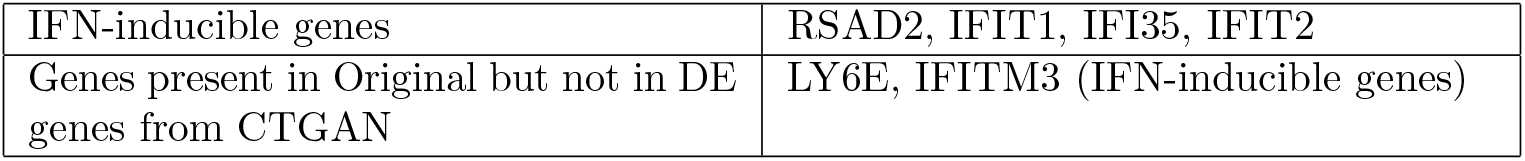
Components associated with extra DE Genes by CTGAN Axes setting with 550 samples.

#### Axes setting

The next logical step is to check how many of the components from Figure 1.e from [26]. are found in the generative models trained on axes genes. GMM was still found to be giving the least number of DE genes even on the largest sample size i.e. 550. The majority of the extra genes still remain associated with the IFN-inducibles component. CTGAN was observed to be missing two IFN-inducible genes which were present in the DE Genes from Original data in Axes setting. CTGAN arguably performs the worst because it only generates extra genes from one component (ISGs) and even then misses 2 genes from the Original data. We also observe the occurrence of STAT1 gene from IFN-γ signalling component in WGAN and GMM. The same was not observed in either of the models in the Non-Axes setting, this could be a direct consequence of reducing the feature space to a biologically significant gene set.

Blankley et.al. also mention that out of all the 16 datasets that were profiled, 5 genes were consistently observed to be differentially expressed in all the datasets. The genes are AIM2, BATF2, FCGR1B, HP and TLR5 (although the exact function of TLR5 in Tuberculosis has not been described in relevant literature), they have been shown to potentially play a role after M. tuberculosis infection [34–37]. Out of the 5 genes, we observe AIM2, BATF2 and FCGR1B to present the list of differentially expressed genes in the original data as well as the generated data from all of the models in both Axes as well as Non Axes settings.

To compare the generated data’s molecular signatures with signatures from other Tuberculosis dataset experiments, we selected a paper by Alam et.al. which talks about the molecular signatures associated with Tuberculosis. The authors have performed overrepresentation analysis for DE Genes obtained from some of the TB datasets present on GEO [25] and came up with a list of biological as well as pathway markers. The authors use PANTHER to perform overrepresentation analysis on the list of DE Genes they accumulated from all the datasets (5,680 in number). The overrepresented ontologies were clustered according to GO Molecular Function, Biological Processes and PANTHER Protein class. The results for this analysis are present as Table 1 in their paper [25].

We performed the same analysis on the DE Genes obtained from our experiments. Over representation analysis was performed(Fisher exact with P-values set according to Bonferroni correction and cutoff set to 0.05). It was observed that very few or no significant results (P-value ¡ 0.05) were obtained with either GO Molecular Functions or GO Cellular components or PANTHER Protein class. Hence, only GO Biological Processes was used to compare the over representation analysis results. We evaluate the models (in both settings) by looking at three aspects as follows:

- Whether there are significant enrichment in the overlapping GO terms.
- Whether there are extra GO terms discovered in generated data which hit with the curated GO terms in the paper [25].
- How many of the GO terms present in the original data set are missed by the generative models.

We report that there is minimal loss of information when reducing feature space of the original data set using Axes Genes. The GO terms over represented by axes genes show a 8/10 overlap with the original data set with 34602 genes (Fig. 7). 16 extra GO terms were observed in the Axes setting and out of these 16 ontology terms, 5 terms show exact overlap with GO terms. Many of the extra ontology were related to each other, a few were “response to cytokines”, “cytokines-mediated signaling pathway”, “response to external stimulus”, “response to stress”, “type I interferon signaling pathway”, etc. Although we also observed 2 GO terms to be missing from Axes data sets compared to non Axes data set, however none of these 2 terms show exact overlap with the table in [25]. This shows that even after reducing the feature space by 99.5%, the biological information suffers minimal losses.

**Figure 7.**
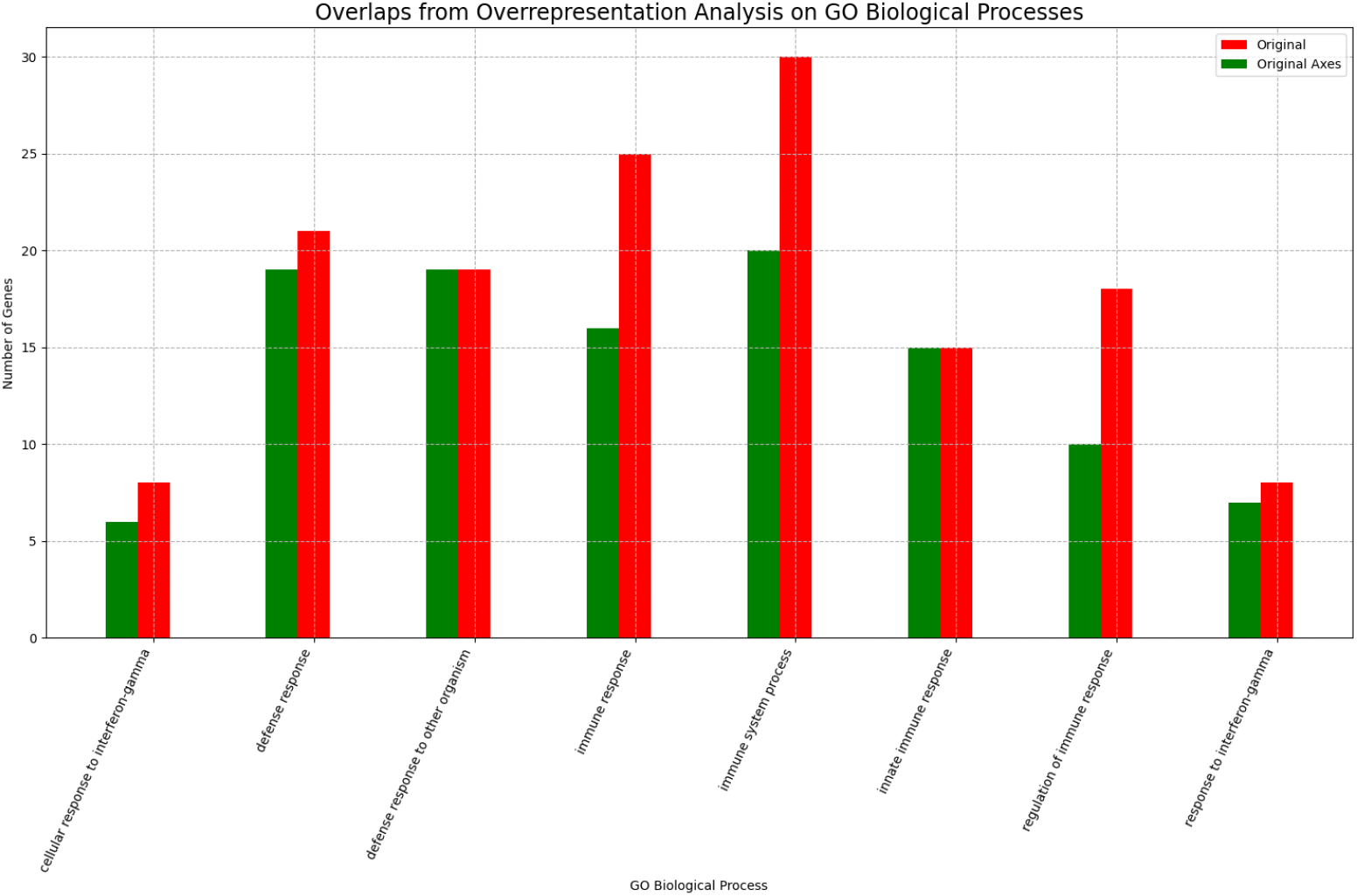
Overlap of Original data (Axes and Non Axes) on GO Biological processes obtained from over representation analysis (P-value<0.05).

### Evaluation of Generative models with GO Biological process

#### WGAN (Non Axes and Axes settings)

In the Non-Axes setting, we observed 50 extra GO Biological Processes terms with 3 exact overlaps with Table 1 from [25]. The GO terms from generated data show a 8/10 overlap with the original data (Fig. 8) with the remaining 2 GO terms not being present in Table 1 from [25]. The enrichment of the common GO terms is also reported in Fig. 8 where the GO terms were enriched by extra genes from the generated data.

**Figure 8.**
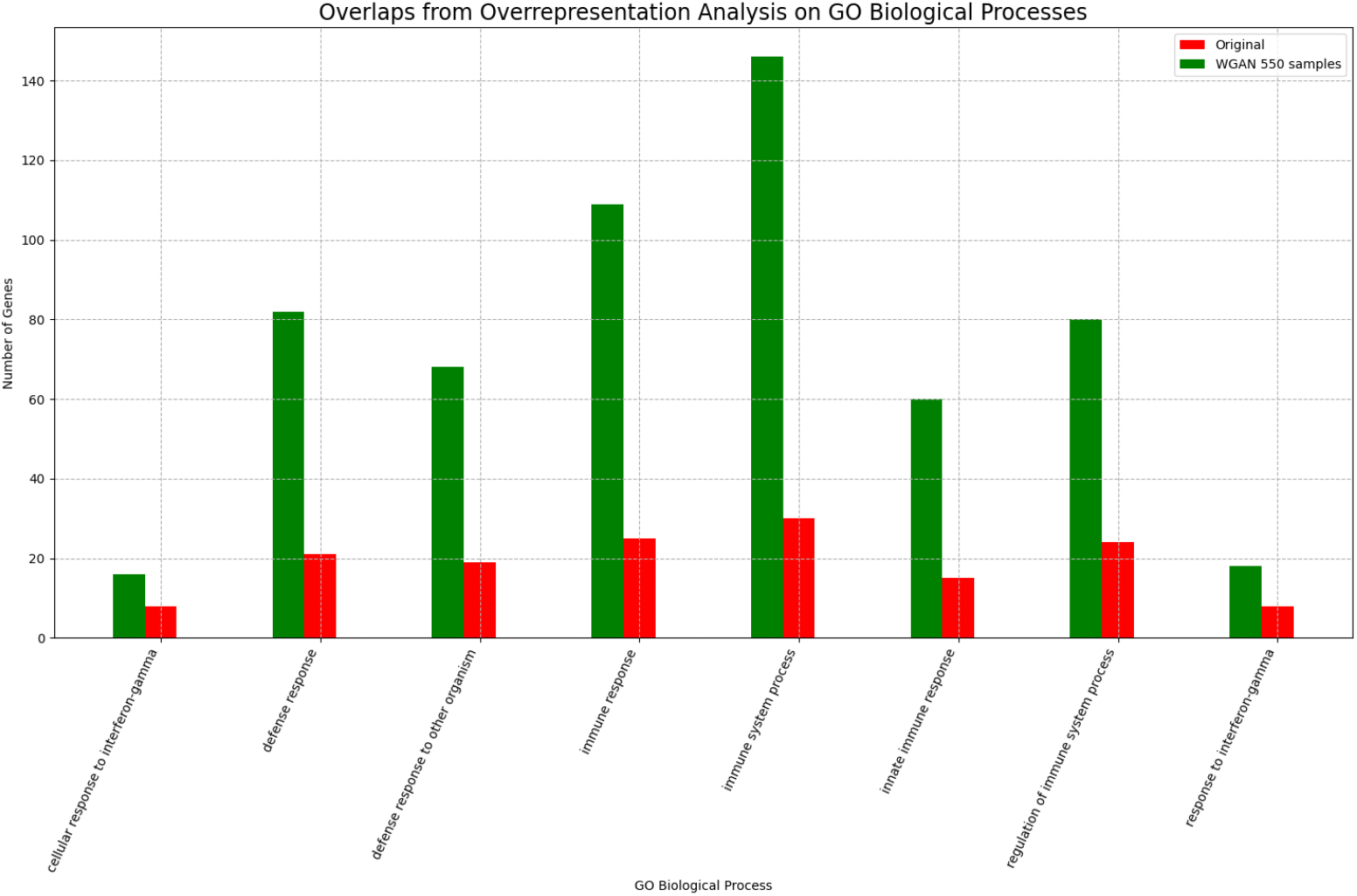
Overlap of WGAN generated data with Original data on GO Biological processes obtained from over representation analysis (P-value<0.05).

A few terms amongst the many related ones were “neutrophil degranulation”, “cellular localization”, “leukocyte activation”, “cellular response to cytokine stimulus”, “interferon-gamma-mediated signaling pathway”, “cell activation involved in immune response”, “antigen processing and presentation of exogenous peptide antigen via MHC class II”, “MHC protein complex assembly “, etc. “Exocytosis” and “Regulation of Exocytosis”, were observed with WGAN in contrast to “Endocytosis” mentioned in Table 1 of [25]. Further, MHC class II and neutrophil degranulation were also captured, which are known mechanistic effects of M. tuberculosis [38, 39]. Presence of such biological processes in the generated data by WGAN might suggest that the model is capturing signals relevant to M.Tuberculosis infection.

Similarly in the Axes setting with WGAN, we observe a 100% (10/10) overlap with the original data under Axes setting with significant enrichments in each GO term (Fig. 9). 46 extra GO terms were reported amongst which none showed exact overlaps. Some of the extra GO terms reported were “interferon-gamma-mediated signaling pathway”, “negative regulation of viral process”, “cellular response to organic substance”, “regulation of response to biotic stimulus”, “antigen processing and presentation of exogenous peptide antigen via MHC class I”, etc.

**Figure 9.**
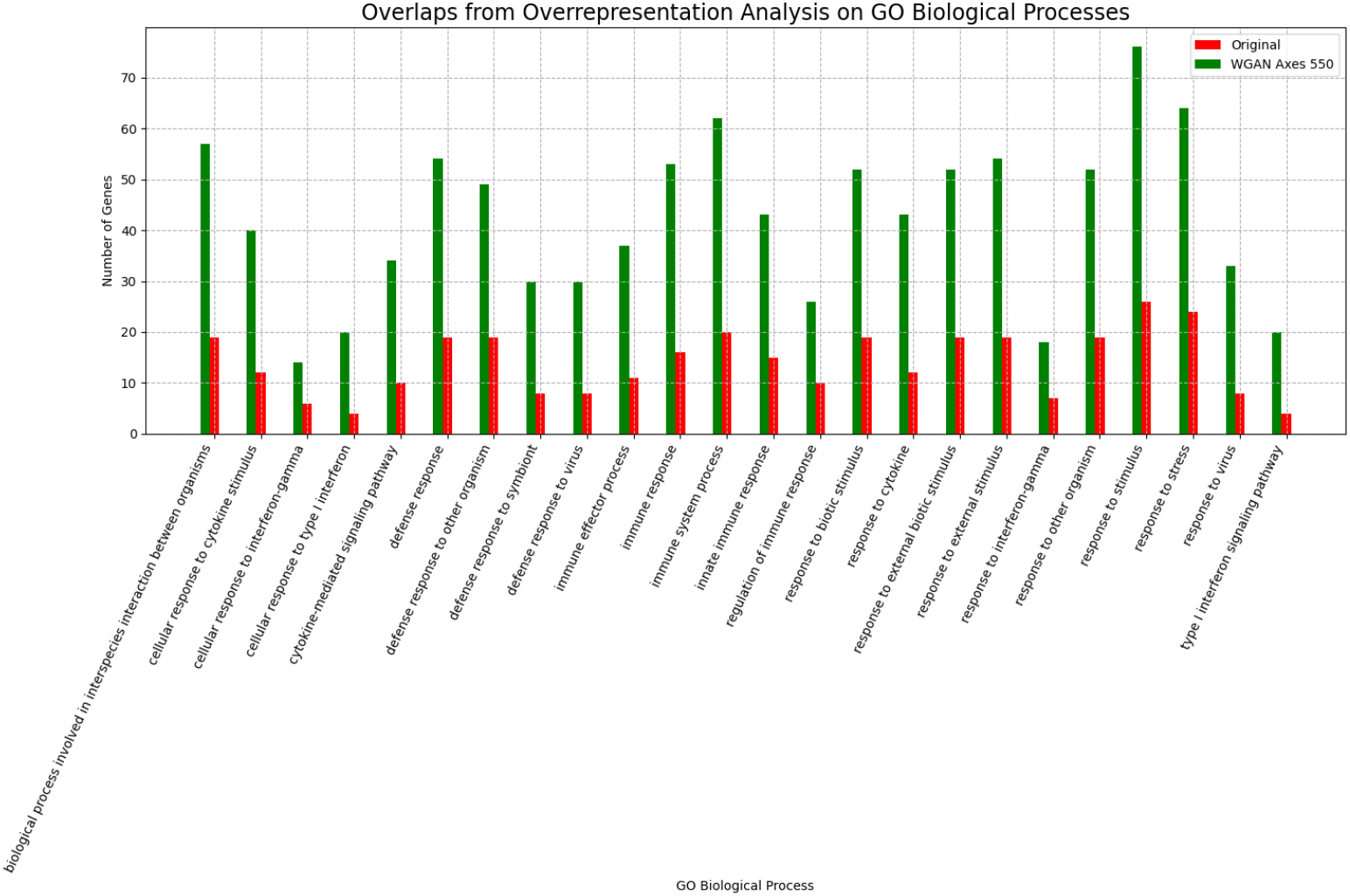
Overlap of WGAN generated data (Axes) with Original data (Axes) on GO Biological processes obtained from over representation analysis (P-value < 0.05).

#### CTGAN (Axes and Non Axes settings)

CTGAN reported a 6/10 overlap (Fig. 10) with original data with the 4 missing terms being related to interferon gamma and regulation of immune response. We report 51 extra GO terms with 6 GO terms overlapping with terms in Table 1 of [25]. Majority of the reported GO terms were related to immunity with “exocytosis” and “Regulation of exocytosis” in agreement with WGAN. The capture of exocytosis as opposed to the previously reported term of endocytosis may indicate the need for carrying out functional experiments to confirm whether the enhanced signal leads to better resolution between these related pathways. As with WGAN, “neutrophil degranulation” and similar GO terms were observed. However, GO terms related to MHC II or MHC I however were not reported in case of CTGAN, possibly indicating superior ability of WGAN to capture biological signal. Under Axes settings, CTGAN reported an 8/10 overlap with original data under Axes settings (Fig. 11), the remaining two being “regulation of immune response” and “response to stimulus”. Enrichments were reported for each GO term, however the enrichments were much weaker when compared to WGAN Axes. There were only 11 extra GO terms reported in case of CTGAN with a majority being related to viral processes and none showing and overlap with Table from [25].

**Figure 10.**
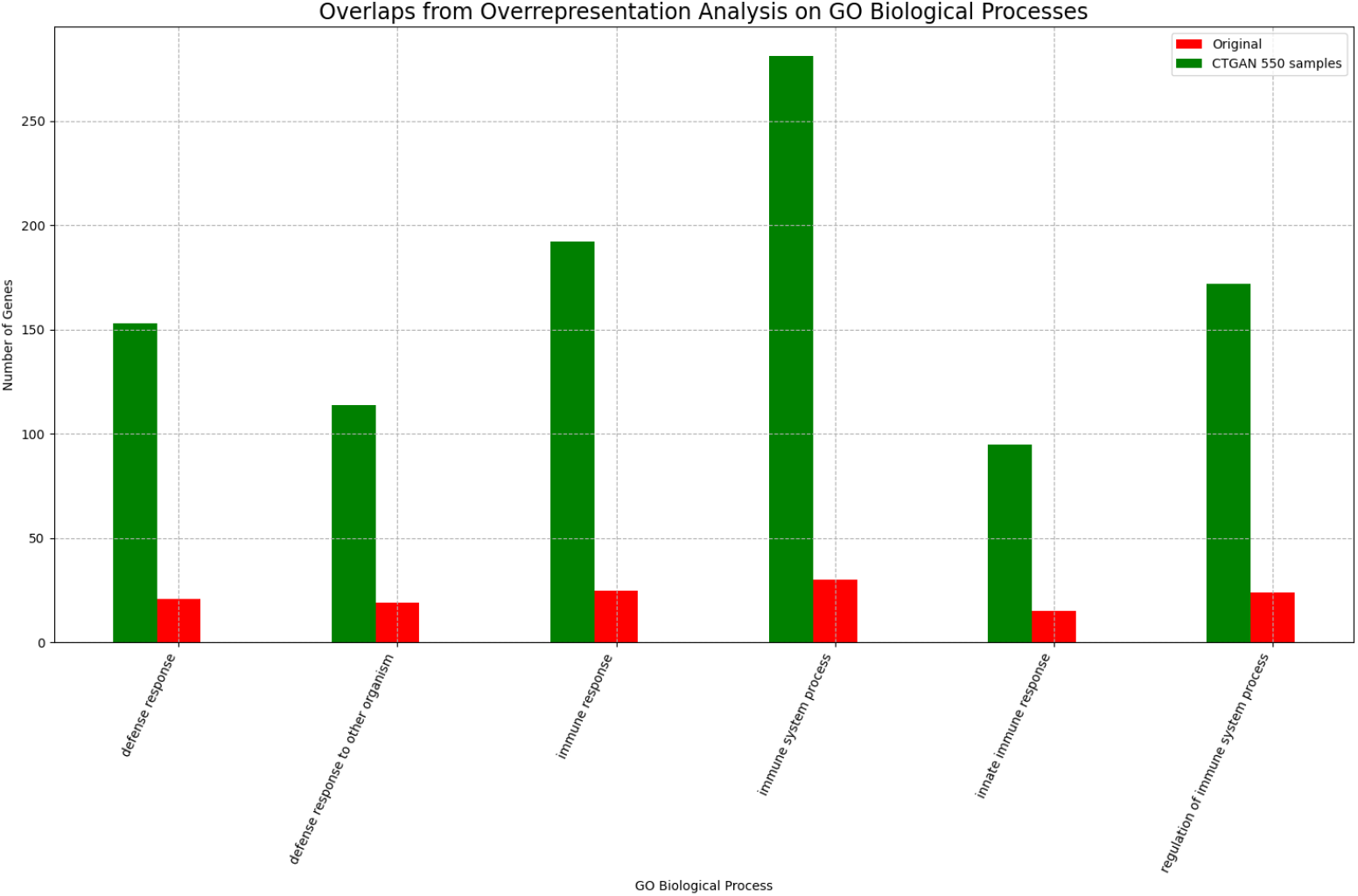
Overlap of CTGAN generated data with Original data on GO Biological processes obtained from overrepresentation analysis (P-value < 0.05).

**Figure 11.**
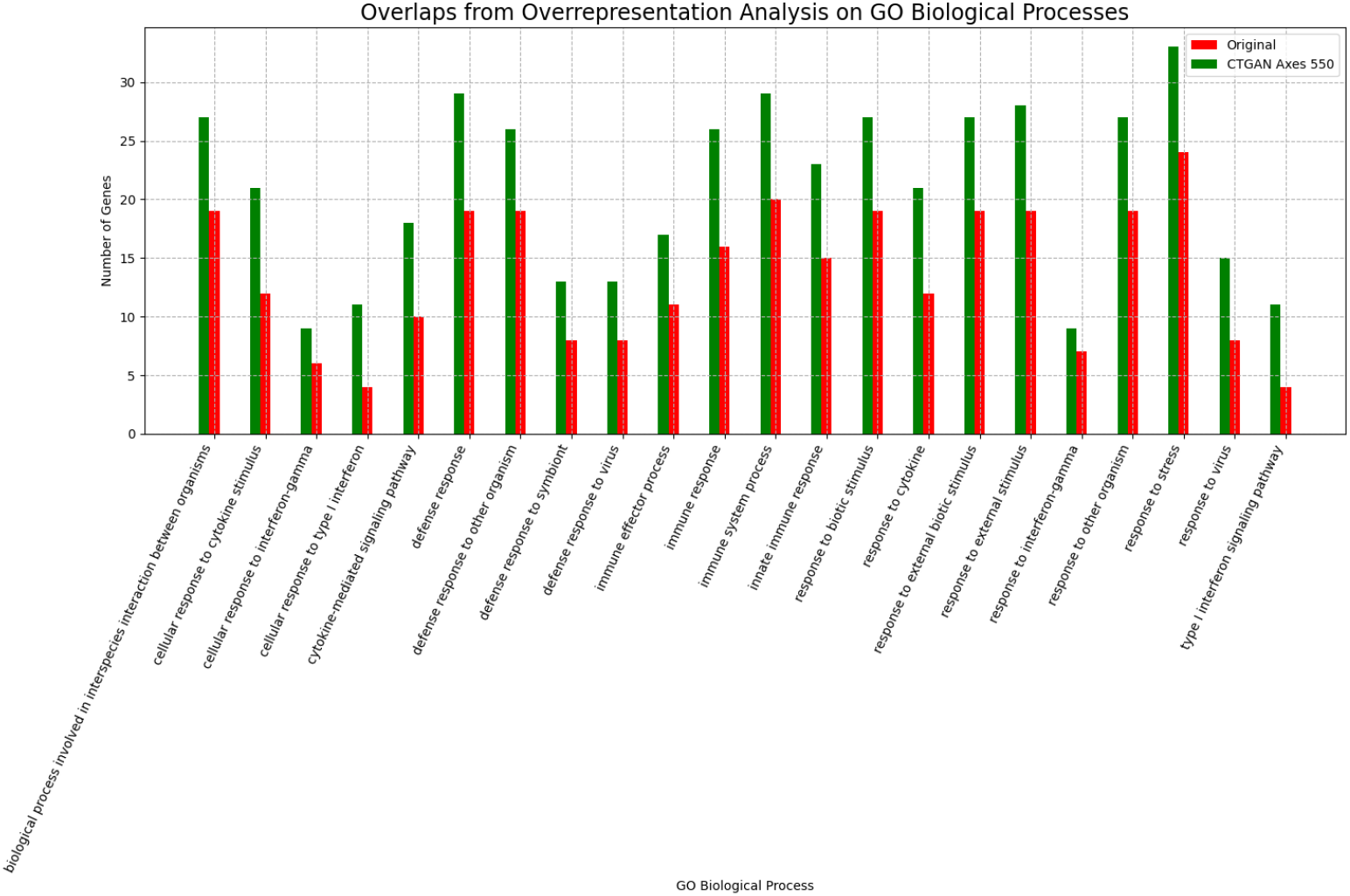
Overlap of CTGAN generated data (Axes) with Original data (Axes) on GO Biological processes obtained from overrepresentation analysis (P-value < 0.05).

#### GMM (Axes and Non Axes settings)

GMM under both the settings showed a 100% (10/10) overlap with the original data, but very little additional signal. This may indicate that GMM is rote-learning from the data without enhancing the signal to noise ratio. Under Non-Axes settings, enrichments were reported for the overlapping ontology terms with original data (Fig. 12), however the enrichments were the weakest when compared to WGAN and CTGAN (under Non-Axes settings). Twenty-two extra GO Terms were reported with 4 showing an exact overlap with the GO Terms in Table 1 from [25]. As expected with a rote-learning method, it reported the known pathways with highest confidence. “Cytokine-mediated signaling pathway”, the most significant GO term from Table 1 from [25], was captured. Interestingly, this pathway was not reported by the WGAN or CTGAN under either settings, perhaps indicating that other components may have a higher enhanced signal than this and would warrant further biological validation. Unlike WGAN and CTGAN, GMM does not report GO terms related to neutrophils or MHCs, and most of the additional captured pathways were related biologically, thus unable to enhance additional signal. Under Axes setting, complete overlap with original (Axes) and 11 extra GO terms were reported. Of these 11 extra terms, only 1 (“cell surface receptor signaling pathway”) showed an overlap with Table 1 from [25], perhaps reinforcing that GMMs are unable to enhance signal from noise and only propagate correlations. Enrichments for overlap with original data are presented in Fig. 13, the enrichments, similar to CTGAN, are weaker than WGAN.

**Figure 12.**
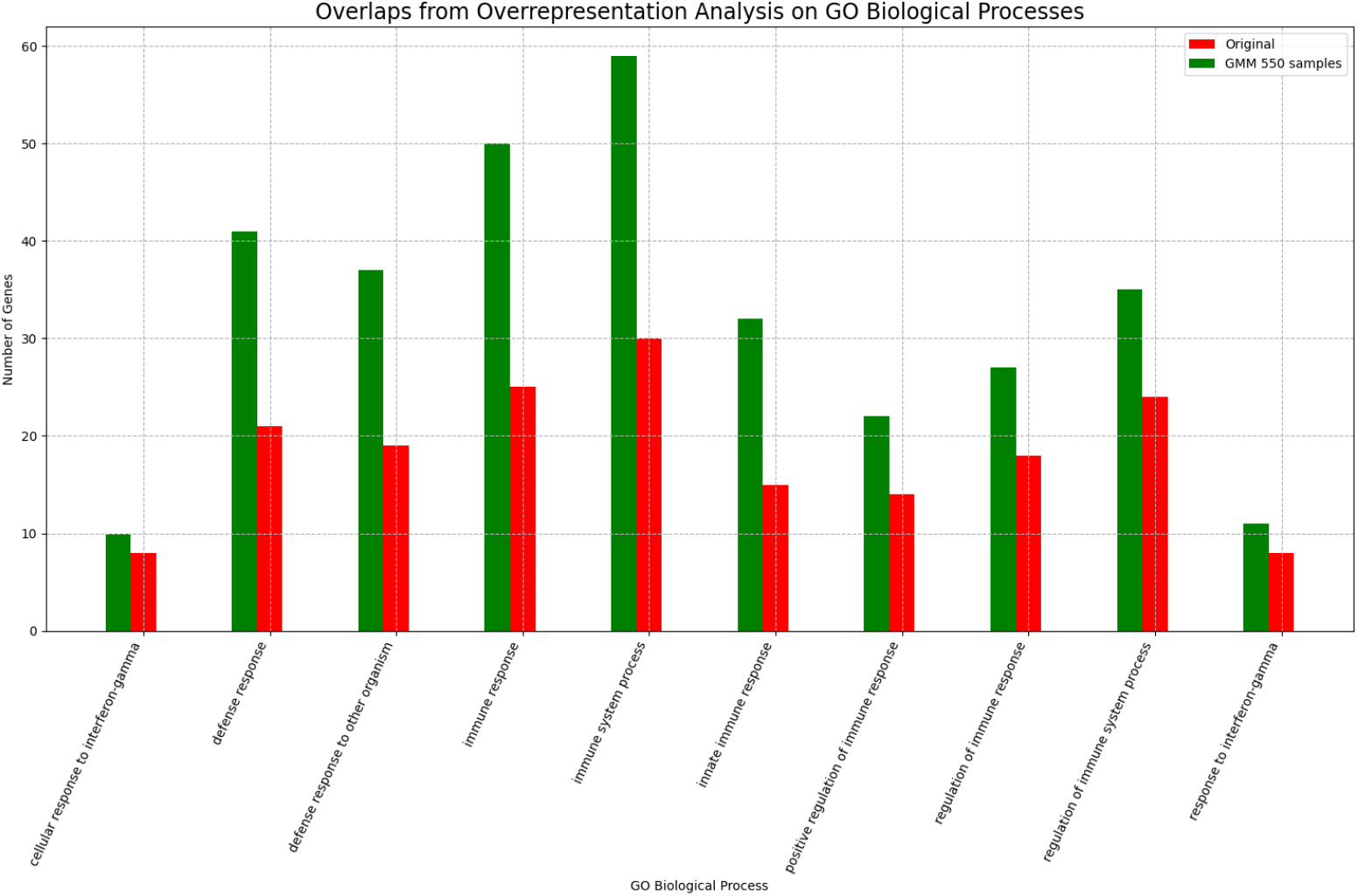
Overlap of CTGAN generated data (Axes) with Original data (Axes) on GO Biological processes obtained from overrepresentation analysis (P-value < 0.05).

**Figure 13.**
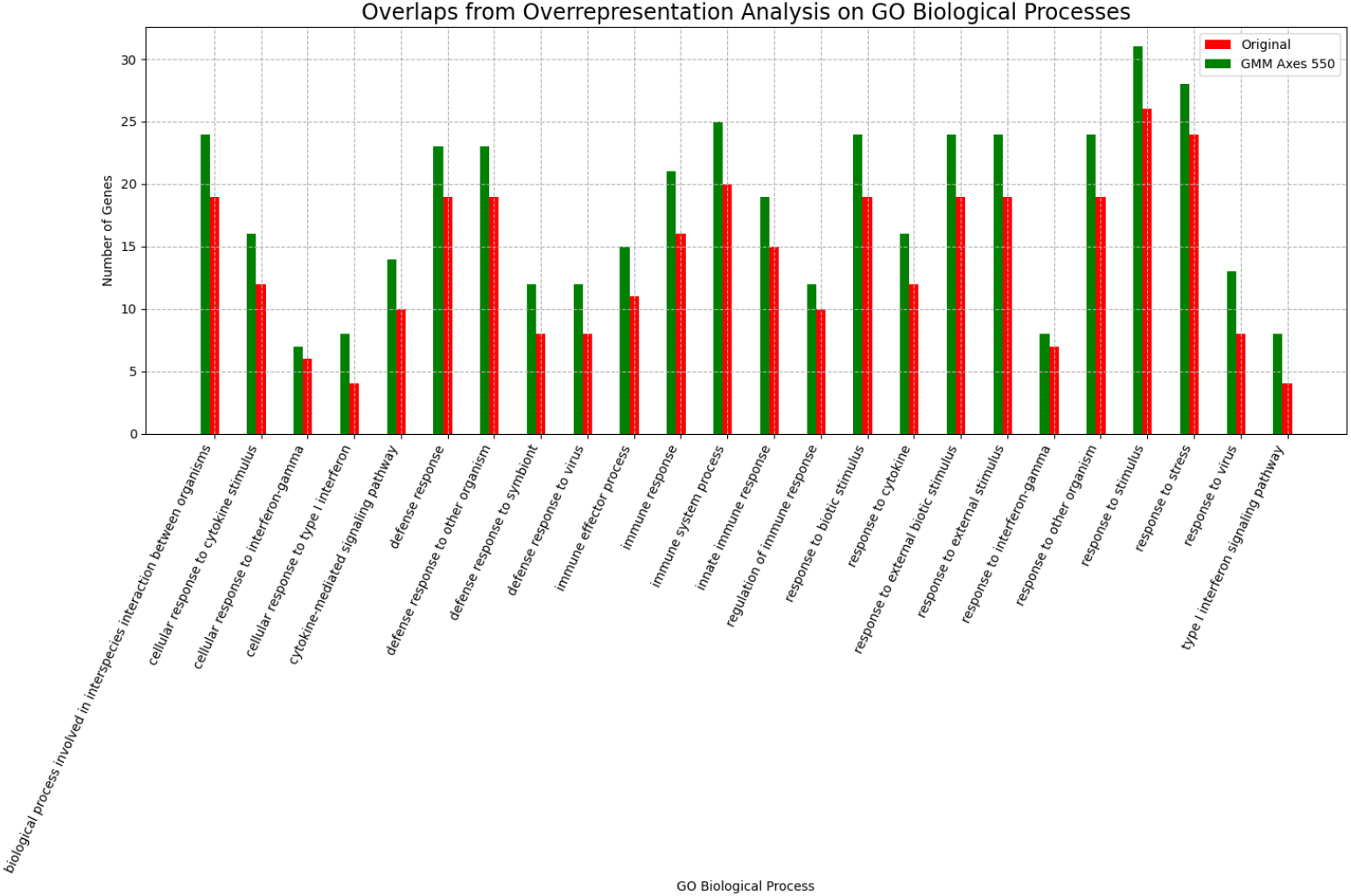
Overlap of CTGAN generated data (Axes) with Original data (Axes) on GO Biological processes obtained from overrepresentation analysis (P-value < 0.05).

## Acknowledgement

TS acknowledges funding support from Department of Biotechnology vide Project BT/PR/34245/AI/133/9/2019.

## Code Availability

https://github.com/tavlab-iiitd/MicroGAN

## Author Contribution

Study Design : TP, Dataset: AG, AS, Modeling: AS, AG, SA, CG, Statistical Analysis: AG, SA, CG, AS, Bioloigical Analysis: SA, CG, AG, TP, RK, Paper writing: SA, AG, CG, TP, RK, AS, HK, Paper Review: TP, RK

## References

1. Draghici S Tarca AL, Romero R. Analysis of microarray experiments of gene expression profiling. American Journal of Obstetrics and Gynecology, 2006.

2. Badr A Zhang G Zhang J, Chiodini R. The impact of next-generation sequencing on genomics. J Genet Genomics, 2011.

3. Bishop DVM et al. Munafò MR, Nosek BA. A manifesto for reproducible science. Nature Human Behaviour, 2017.

4. Mokrysz C et al. Button KS, Ioannidis JPA. Power failure: Why small sample size undermines the reliability of neuroscience. Nature Reviews Neuroscience, 2013.

5. Klein U ZTu Y, Stolovitzky G. Quantitative noise analysis for gene expression microarray experiments. Proceedings of the National Academy of Sciences of the United States of America., 2002.

6. Meng F. Scherer A, Dai M. Impact of experimental noise and annotation imprecision on data quality in microarray experiments. Methods in Molecular Biology., 2013.

7. Hall LO et al. Chawla N v., Bowyer KW. Smote: Synthetic minority over-sampling technique. Journal of Artificial Intelligence Research, 2002.

8. Mirza M et al Goodfellow IJ, Pouget-Abadie J. Generative adversarial nets. Advances in Neural Information Processing Systems., 2014.

9. Probabilistic Graphical Models. The MIT Press.

10. Williams C et al Beaulieu-Jones BK, Wu ZS. Privacy-preserving generative deep neural networks support clinical data sharing. Circulation: Cardiovascular Quality and Outcomes., 2019.

11. Lenz S et al. Nußberger J, Boesel F. Synthetic observations from deep generative models and binary omics data with limited sample size. Briefings in Bioinformatics., 2020.

12. Kotecha K. Chaudhari P, Agrawal H. Data augmentation using mg-gan for improved cancer classification on gene expression data. Soft Computing., 2020.

13. Greene CS. Way GP. Extracting a biologically relevant latent space from cancer transcriptomes with variational autoencoders. Pacific Symposium on Biocomputing., 2018.

14. Bansal V et al. Marouf M, Machart P. Realistic in silico generation and augmentation of single-cell rna-seq data using generative adversarial networks. Nature Communications., 2020.

15. Huang H. Wang X, Ghasedi Dizaji K. Conditional generative adversarial network for gene expression inference. Bioinformatics., 2018.

16. Smirnov P et al. Rampášek L, Hidru D. Dr.vae: improving drug response prediction via modeling of drug perturbation effects. Bioinformatics, 2019.

17. Russell C et al. Srivastava A, Valkov L. Veegan: Reducing mode collapse in gans using implicit variational learning. Advances in Neural Information Processing Systems., 2017.

18. Pfau D et al. Metz L, Poole B. Unrolled generative adversarial networks. ICLR conference papers., 2016.

19. Taylor G et al. Im DJ, Ma H. Quantitatively evaluating gans with divergences proposed for training. arXiv., 2018.

20. Google MM et al. Lucic M, Kurach K. Are gans created equal? a large-scale study. Advances in Neural Information Processing Systems., 2018.

21. Weiss Y. Richardson E. On gans and gmms. Advances in Neural Information Processing Systems., 2018.

22. Menius A et al. Su Y, Zhu L. Mixture models for gene expression experiments with two species. Human Genomics., 2014.

23. Poehlman WL et al. Ficklin SP, Dunwoodie LJ. Discovering condition-specific gene co-expression patterns using gaussian mixture models: A cancer case study. Scientific Reports., 2017.

24. Kim J et al. Preininger M, Arafat D. Blood-informative transcripts define nine common axes of peripheral blood gene expression. PLoS Genetics., 2013.

25. Ahmed MM et al. Alam A, Imam N. Identification and classification of differentially expressed genes and network meta-analysis reveals potential molecular signatures associated with tuberculosis. Frontiers in Genetics., 2019.

26. Levin J et al. Blankley S, Graham CM. A 380-gene meta-signature of active tuberculosis compared with healthy controls. European Respiratory Journal., 2016.

27. Oni T et al. Kaforou M, Wright VJ. Detection of tuberculosis in hiv-infected and -uninfected african adults using whole blood rna expression signatures: A case-control study. PLoS Medicine., 2013.

28. Hinds J et al. Muttucumaru DGN, Roberts G. Gene expression profile of mycobacterium tuberculosis in a non-replicating state. Tuberculosis., 2004.

29. Fitzgerald KA. The interferon inducible gene: Viperin. Journal of Interferon and Cytokine Research., 2011.

30. McNab FW et al. O’Garra A, Redford PS. The immune response in tuberculosis. Annual Review of Immunology., 2013.

31. Schlesinger LS. Mycobacterium tuberculosis and the complement system. Trends in Microbiology., 1998.

32. Tripathy SP et al. Hilda JN, Das S. Role of neutrophils in tuberculosis: A bird’s eye view. Innate Immunity., 2020.

33. Galli L et al. de Martino M, Lodi L. Immune response to mycobacterium tuberculosis: A narrative review. Frontiers in Pediatrics., 2019.

34. Shimada Y et al. Saiga H, Kitada S. Critical role of aim2 in mycobacterium tuberculosis infection. International Immunology., 2012.

35. Murphy TL et al. Tussiwand R, Lee WL. Compensatory dendritic cell development mediated by batf-irf interactions. Nature., 2012.

36. Gutschmidt A et al. Jacobsen M, Repsilber D. Candidate biomarkers for discrimination between infection and disease caused by mycobacterium tuberculosis. Journal of Molecular Medicine., 2007.

37. Mvundura E et al. Kasvosve I, Gomo ZAR. Haptoglobin polymorphism and mortality in patients with tuberculosis. International Journal of Tuberculosis and Lung Disease, 2000.

38. Kinnear C et al. Kroon EE, Coussens AK. Neutrophils: Innate effectors of tb resistance? Frontiers in Immunology., 2018.

39. Rojas RE et al. Torres M, Ramachandra L. Role of phagosomes and major histocompatibility complex class ii (mhc-ii) compartment in mhc-ii antigen processing of mycobacterium tuberculosis in human macrophages. Infection and Immunity., 2006.

